# Genetic heterogeneity in the *Salmonella* Typhi Vi capsule locus: A population genomic study from Fiji

**DOI:** 10.1101/2023.12.04.569801

**Authors:** Aneley Getahun Strobel, Andrew J. Hayes, Wytamma Wirth, Mikaele Mua, Tiko Saumalua, Orisi Cabenatabua, Vika Soqo, Varanisese Rosa, Nancy Wang, Jake A. Lacey, Dianna Hocking, Mary Valcanis, Adam Jenney, Benjamin P. Howden, Sebastian Duchene, Kim Mulholland, Richard A. Strugnell, Mark R. Davies

## Abstract

Typhoid fever is endemic in many parts of the world and remains a major public health concern in tropical and sub-tropical developing nations, including Fiji. To address high rates of typhoid fever, the Northern Division of Fiji is implementing a mass vaccination with typhoid conjugate vaccine (Vi-polysaccharide conjugated to tetanus toxoid) as a public health control measure in 2023. In this study we define the genomic epidemiology of *S*. Typhi in the Northern Division prior to island-wide vaccination, sequencing 85% (n=419) of the total cases from the Northern Division and Central Divisions of Fiji that occurred in the period 2017-2019. We found elevated rates of nucleotide polymorphisms in *tviD and tviE* genes (responsible for Vi-polysaccharide synthesis) relative to core genome levels within the Fiji endemic *S*. Typhi genotype 4.2. Expansion of these findings within a globally representative database of 12,382 *S*. Typhi (86 genotyphi clusters) showed evidence of convergent evolution of the same *tviE* mutations across the *S*. Typhi population, indicating that *tvi* selection has occurred both independently and globally. The functional impact of *tvi* mutations on the Vi-capsular structure and other phenotypic characteristics are presently unknown, yet commonly occurring *tviE* polymorphisms localise adjacent to predicted active site residues when overlayed against the predicted TviE protein structure. Given the central role of the Vi-polysaccharide in *S*. Typhi biology and vaccination, further integrated epidemiological, genomic, and phenotypic surveillance is required to determine the spread and functional implications of these mutations.

## Introduction

Typhoid fever is a major public health concern worldwide and is caused by the bacterium *Salmonella enterica* subsp. *enterica* serovar Typhi (*S*. Typhi). In 2017, it had an estimated global incidence of over 10 million cases per year leading to over 100,000 deaths^1^. Fiji is an island nation in the South Pacific with the population of 884,887^2^ where typhoid fever remains endemic^3,4^. The reported incidence of *S.* Typhi infections in Fiji has risen over the last two decades, with the number and incidence of culture-confirmed typhoid cases in Fiji increasing from <5/100,000^5^ to over 40/100,000 in the mid 2000s^6,7^. This apparent increase in typhoid fever may have been linked with the establishment of laboratory surveillance in 2004 and 2005 which could have resulted in more systematic reporting of laboratory confirmed cases and improved diagnostic facilities^8^. In addition, population growth in peri-urban locations, poor sanitation, hygiene, and inadequate water supply could have contributed to the increased transmission and reporting of typhoid in rural and peri-urban settlement^8,9^.

A previous study of typhoid cases in the Central Division of the island of Viti Levu, Fiji, showed that similar to other Pacific Island Countries^10,11^, the burden of disease is largely due to local endemic strains. In the Central Division of Fiji, the archetypical *S*. Typhi strain is defined as genotype 4.2, with sporadic cases related to the globally dominant genotype 4.3.1^12^. Genotype 4.2 strains showed low rates of evolution similar to other well described typhoid lineages; however, genotype 4.2 populations lack genetic^12^ or phenotypic^5–7^ resistance to antimicrobials that is a hallmark of the globally dominant genotype 4.3.1 (H58) population^13^.

Three types of commercially available vaccines are utilised worldwide to combat typhoid. These vaccines have an efficacy of between 50-96% within the first year with diminishing returns in subsequent years^14^. The vaccines include: a live-attenuated *S*. Typhi strain Ty21a which does not express the Vi capsular polysaccharide; Vi polysaccharide-based (e.g. Typhim Vi and Vivaxim) and, more recently, new protein-conjugated Vi (e.g. Typbar-TCV and TYPHIBEV). The proteins that assemble, modify and secrete Vi polysaccharide in *S*. Typhi are encoded within a 10-gene operon within the *viaB* locus of *Salmonella* pathogenicity island 7 (SPI-7) and the functional operon is important for *S*. Typhi virulence^15,16^. The operon consists of 4 genes that encode enzymes for Vi biosynthesis (*tvi* genes) and 5 involved in translocation (*vex* genes). The gene *tviA* is the first in the operon and acts as the transcriptional regulator of the operon. The genes *tviB* and *tviC* produce the monomeric sugar nucleoside unit (UDP-GalNAcA) that is polymerised by the GT4 family glycosyltransferase encoded by *tviE* while the TviD protein was recently found to act as an O-acetyltransferase that modifies the polymer through acetylation^17^. Mutations in the *viaB* operon have been shown to be overrepresented in chronic carriage isolates in the gall bladder^18^ and *tvi* genes were identified to carry a higher level of mutations in a 2008 study that examined 19 isolates of *S.* Typhi^19^; however no comprehensive study of *tvi* mutations in a global genomics dataset with global prevalence, has been performed to-date.

Historically, rates of typhoid fever have been higher in the Northern Division than in the Central Division of Fiji^6,7,8^. As part of the public health response, the Fijian Ministry of Health has utilised vaccination to curb the spread of typhoid outbreaks in local health jurisdictions^6^, and in July 2023, initiated a mass vaccination with Typbar-TCV (Vi polysaccharide conjugated to a tetanus toxoid) in the Northern Division^20^. In this study, we defined the population structure of *S*. Typhi strains in the Northern Division between 2017-2019, i.e. prior to the commencement of the mass vaccination campaign. We also undertook a comparative phylogenetic investigation of *S*. Typhi in the Northern Division relative to the Central Division of Fiji to better define the spread of typhoid clones in Fiji, and whether there was likely to be frequent between-island transmission of the pathogen. Considering the importance of Vi in *S*. Typhi biology and disease control, we conducted a regional and international analyses of the Vi biosynthesis genes, to define the underlying genomic landscape of Vi evolutionary dynamics.

## Methods

### Study setting and sites

The primary study site is the Northern Division which is comprised of the second largest island in Fiji, Vanua Levu, and other smaller islands. It has a total population of 131,918 which accounts for 15% of Fiji’s total population^2^. It is further divided into four sub-divisions: Bua, Cakaudrove, Macauta and Taveuni for health services administration. The Central Division, comprising about half of the main Fijian island of Viti Levu, is the secondary site for the comparative genomic analysis. It has a population of 378,284 representing 42.7% of the total Fijian population.

### Study participants identification and data collection

All culture-confirmed typhoid fever patients from the Northern and Central Divisions between 1 January 2017 and 31 December 2019 were eligible for inclusion in this study. A standardised case investigation form was used to collect demography, and epidemiological status including travel history, outbreaks, attendance of social gathering etc. Data was collected through direct interviews, from health facility medical records, laboratory registers, health inspectors and infection control nurse reports.

In Fiji, the outbreak response for typhoid fever is triggered by the sudden increase in the number of typhoid cases (compared with the previous reporting period) or the report of two or more patients with typhoid, in the same household or community, in a four-week period^21^. For this study, epidemiologically-linked outbreak clusters were defined as two or more typhoid patients from the same household (household clusters) including visitors and relatives, or from the same community village/neighbourhood (community clusters), as determined during case or outbreak investigation or study participant interviews.

### Bacterial culture and antimicrobial susceptibility testing

All bacterial cultures were performed in two microbiology laboratories in Fiji (Labasa Hospital in the Northern and Colonial War Memorial Hospital in the Central Division) following the local standard operating procedure. Briefly, blood cultures were processed using a BaCT/ALERT 3D (BioMerieux, Marcy L’Etoile, France) system. Blood cultures with Gram-negative bacilli were inoculated onto Sheep Blood, Chocolate, MacConkey (MC) agar plates (Difco, Maryland, USA) and Triple Sugar Iron agar (Difco, Maryland, USA). Blood agar and Chocolate agar plates were incubated in CO₂ using candle jar for 18 to 24 hours. While MC and TSI were incubated for 18 to 24 hours at 37°C aerobically. Stool samples were first inoculated into Selenite (Sel) broth (Difco, Maryland, USA) (approximately 1 g of stool in 10 ml of Sel broth) and incubated overnight at 37°C before inoculation onto MC, Xylose Lysine Deoxycholate (Difco, Maryland, USA) and *Salmonella* chromogenic agar plates (Difco, Maryland, USA). The growth of non-lactose fermenting organisms on MC agar (colourless and transparent colonies) and the presence of traces of hydrogen sulphide which appeared as black layer at the base of the TSI slant and glucose fermented base with the absence of gas production (K/A+) suggested the growth of *S.* Typhi. A positive agglutination reaction with Polyvalent O, D, O9, and Vi antisera confirmed the growth of *S.* Typhi.

The antimicrobial susceptibility test was performed using Disk Diffusion method on Mueller-Hinton agar (Difco, Maryland, USA). Antimicrobials routinely tested included ampicillin (10 µg), cephalothin (30 µg), ceftriaxone (30 µg), chloramphenicol (30 µg), gentamicin (10 µg), sulfamethoxazole (23.75 µg) + trimethoprim (1.25 µg), ciprofloxacin (5 µg) and nalidixic acid (30 µg). Interpretation of susceptibility tests was conducted using the Clinical and Laboratory Standard Institute guidelines.

### Whole genome sequencing, genotyping and *in silico* detection of AMR genes

Whole genome sequencing (WGS) was conducted at the Microbiological Diagnostic Unit Public Health Laboratory (MDU PHL) at the University of Melbourne, Australia. DNA was extracted using a QIAsymphony DSP virus pathogen kit (Qiagen, Maryland, USA). Genome sequencing was performed using Illumina Nextera XT (Illumina, California, USA) on the Illumina NextSeq 500/550 platform using 150 paired-end reads. In total, 419 *S*. Typhi strains (240 from Northern and 179 from Central Division) were sequenced. *De novo* draft genome assemblies were generated from Illumina short reads using SPAdes v v3.14.1 and were screened for known antimicrobial resistance determinants in AMRFinderPlus (https://github.com/MDU-PHL/abritamr) using abriTAMR (v0.9.8) with a minimum nucleotide identity of 90% and minimum coverage of 100%. Genotyping of the genome sequences was determined using the genotyphi framework (https://github.com/katholt/genotyphi)^22^.

### Mapping and SNP calling

To determine phylogenetic structure in the Fijian 4.2.1 and 4.2.2 subclades, 782 genomes (419 from this study and 363 previously published^12,22^) from these clades were mapped to the Fijian *S.* Typhi reference strain ERL072973 (genotype 4.2.2, accession number LT904777.2). Illumina paired-end short reads were mapped to ERL072973 using bwa-mem2 as part of snippy v4.6.0 (github/tseemann/snippy). A total of 1265 SNP and indel events were identified in the mapping using FreeBayes v1.3.1^23^ and functional annotations of SNPs called using SnpEff v4.3t^24^ as part of snippy^25^. For core genome determination and tree building, mobile genetic elements and genomic regions of irregular SNP density were identified in the reference genome and the isolate core genome alignment using Gubbins v[2.4.1]^26^. All low complexity mapping regions, high SNP density regions and mobile genetic elements were then excised from the alignment resulting in a 4,569,951bp core genome alignment consisting of a total of 828 core genome SNPs, including 326 parsimony informative sites and 502 singleton sites.

### Phylogenetic analysis

Consensus SNP alignments were used to build a maximum-likelihood tree with IQ-TREE v1.6.12^27^. A general time-reversible model with gamma distribution for among-site rate heterogeneity (GTR+F+G4) was used in IQ-TREE, performed with 1000 non parametric bootstrap replicates to assess toplogical uncertainty^28^. Pairwise SNP distance between each *S.* Typhi isolate was calculated from core genome SNPs using the dist.dna function in *ape* (model=”N”) in R^29^.

### Phylodynamic analysis

Individual phylodynamic analyses were performed on each lineage (4.2.1 and 4.2.2) because they are assumed to behave like independent populations. All phylodynamic analyses were performed on core SNP alignments, with the number of constant sites specified, in Bayesian evolutionary analysis by sampling trees (BEAST) v1.10.4^30^. The midyear date was used for isolates where the precise date of isolation was not available. Bayesian Evaluation of Temporal Signal (BETS) was used to test for temporal signal and to select a molecular clock model in a fully Bayesian context^31^. The effective population size of each lineage was calculated using separate Bayesian Skygrid tree prior for each lineage^32^. Geographic Divisions (Central and Northern) and four subdivisions; Bua, Cakaudrove, Macuata and Taveuni in the Northern Division were treated as discrete traits in a Bayesian phylogeographic model, that directly estimates the number of migration events between locations^33^. Maximum–clade credibility (MCC) phylogenetic trees were constructed for each lineage to visualise the evolutionary history of isolates. Additional metadata from this study and case control study^12^ was used to assess the relationship between isolates and transmission patterns. Markov chain Monte Carlo (MCMC) was used to sample the posterior distribution, as implemented in BEAST, with sufficient sampling determined according to an effective sample size of 200 for key model parameters, as visualised in Beastiary^34^). Importantly, recent research suggests that discrete phylogeographic analyses are sensitive to the choice of prior distributions on the average dispersal rate and the number of dispersal routes. We conducted prior sensitivity analyses, as recommended by Gao et al.^35^, which suggested that our results here are robust to the choice of the default priors.

### Mutational analysis within a global *S*. Typhi database

To screen for evidence of mutational selection in the *viaB* operon, and elsewhere in the core genome, the SNP profile of the Fijian genotype 4.2 population was compared with a global dataset of 12,382 *S.* Typhi genomes constituting 86 genotyphi clusters^13^. Single nucleotide polymorphisms (SNPs) were called by mapping of assembled sequences to the CT18 reference strain (NC_003198.1) using minimap2 (v2.18)^36^. Individual variant call files (VCF) were merged with bcftools (v1.14) yielding a total of 124,180 unique SNPs. Further filtering of the VCF was performed to remove events present in only a single isolate, reducing the number of SNPs to 36,432. Once variants were called functional effects of variants was called with snpEff v4.3^24^ using annotations from the genbank CT18 reference genome (NC_003198.1). Analysis of gene mutation profiles relative to individual genotyphi clusters was performed in R v4.1.1 using the following packages; gggenes v0.4.1, see v0.7.0, ggrepel v0.9.1, dplyr v1.0.8, tidyr v1.2.0 and ggplot2 v3.3.5. A ratio of non-synonymous to synonymous mutations in a gene was calculated for visualisation on the plots. Where no synonymous mutations were present in a gene the value was assumed to be the maximum possible by adding 1 to the synonymous count to allow calculation even when no synonymous mutations were present in a gene. The overall low rates of synonymous mutations in the dataset prevented a model-informed dn/ds ratio to be inferred^37^.

### Mapping of mutational hotspots onto TviE protein structure

To determine if the mutations over-represented in Fiji and the global dataset occurred in similar locations on the 3D structure of the TviE protein, mutations were mapped to an available structure. An Alphafold predicted structure was obtained from Uniprot (https://www.uniprot.org/uniprotkb/Q04975/entry, https://alphafold.ebi.ac.uk/entry/Q04975, AF-Q04975-F1-model_v2.pdb). Model confidence was very high (pLDDT > 90) for the majority of residues, suggesting the predicted structure is an appropriate analogue for the protein structure. UCSF chimera was used to visualise the mutations against the predicted protein structure^38^. As a proxy for independent acquisition of mutations, presence in multiple genotyphi clusters was plotted. Residues with mutations in > 5 genotyphi clusters (>5% of total clusters, 95% quantile of distribution of clusters) were coloured in blue, while residues with mutations in > 10 genotyphi clusters (>10% of total clusters) were coloured in red. The structure for TviD (https://www.uniprot.org/uniprotkb/Q04974/entry) was also assessed to determine the feasibility of plotting mutations on the structure however only about half the residues had very high confidence, with several residues of Low confidence prediction, suggesting the model may have inaccuracies.

### Ethics

The study received ethics approval from Fiji National Health Research Ethics Review Committee (ID number: 2016.86 and 2018.132) and the human research ethics committee of the University of Melbourne (ID number: 1852159).

### Data availability

Illumina sequence reads were deposited into the European Nucleotide Archive (Bioproject identifier PRJNA1032150). Accession numbers for sequence reads are supplied in Supplementary table 1.

## Results

### Demographic characteristics of Northern Division typhoid fever patients

A total of 246 culture-confirmed typhoid fever patients were reported in the Northern Division during 2017-2019. Of these, 218 were blood culture confirmed, 19 blood and stool, eight stool and one abscess fluid culture confirmed. The majority of patients (93.8%) were from the iTaukei ethnic group, 58.1% were males and 83% resided in rural areas. The median age was 26 years (IQR 17-41). Children (<15 year of age) represented 22% of the overall patients (Table 1).

**Table 1.** Demographic characteristics of culture-confirmed typhoid fever patients in the Northern Division (N=241*)

| Characteristics | Culture-confirmed cases. Total number (%) |
| --- | --- |
| **Year** |  |
| 2017 | 113 (46.9) |
| 2018 | 64 (26.6) |
| 2019 | 64 (26.6) |
| **Gender** |  |
| Male | 140 (58.1) |
| Female | 101 (41.9) |
| **Ethnicity** |  |
| iTaukei | 226 (93.8) |
| Fijian of Indian Descent | 8 (3.3) |
| Other ethnicity | 7 (2.9) |
| **Age** |  |
| <15 years | 53 (22) |
| ≥15 years | 188 (78) |
| **Location** |  |
| Rural | 200 (83) |
| Urban | 41 (17) |
| **Subdivisions** |  |
| Cakaudrove | 118 (49) |
| Macuata | 76 (31.5) |
| Taveuni | 26 (10.8) |
| Bua | 21 (8.7) |

The average incidence of typhoid fever was 60.9 cases/100,000 population (95% CI 47.6, 74.2). The average incidence of typhoid fever increased progressively with age and reached its peak in the 20-29 year old age groups. High incidence (>100 cases/100,000 population) was reported in individuals between 20-34 year old age groups among males and 20-29 year old age groups in women. A second smaller peak in incidence was observed between the age of 40-49 years for both females and males. (Figure 1).

**Figure 1.**
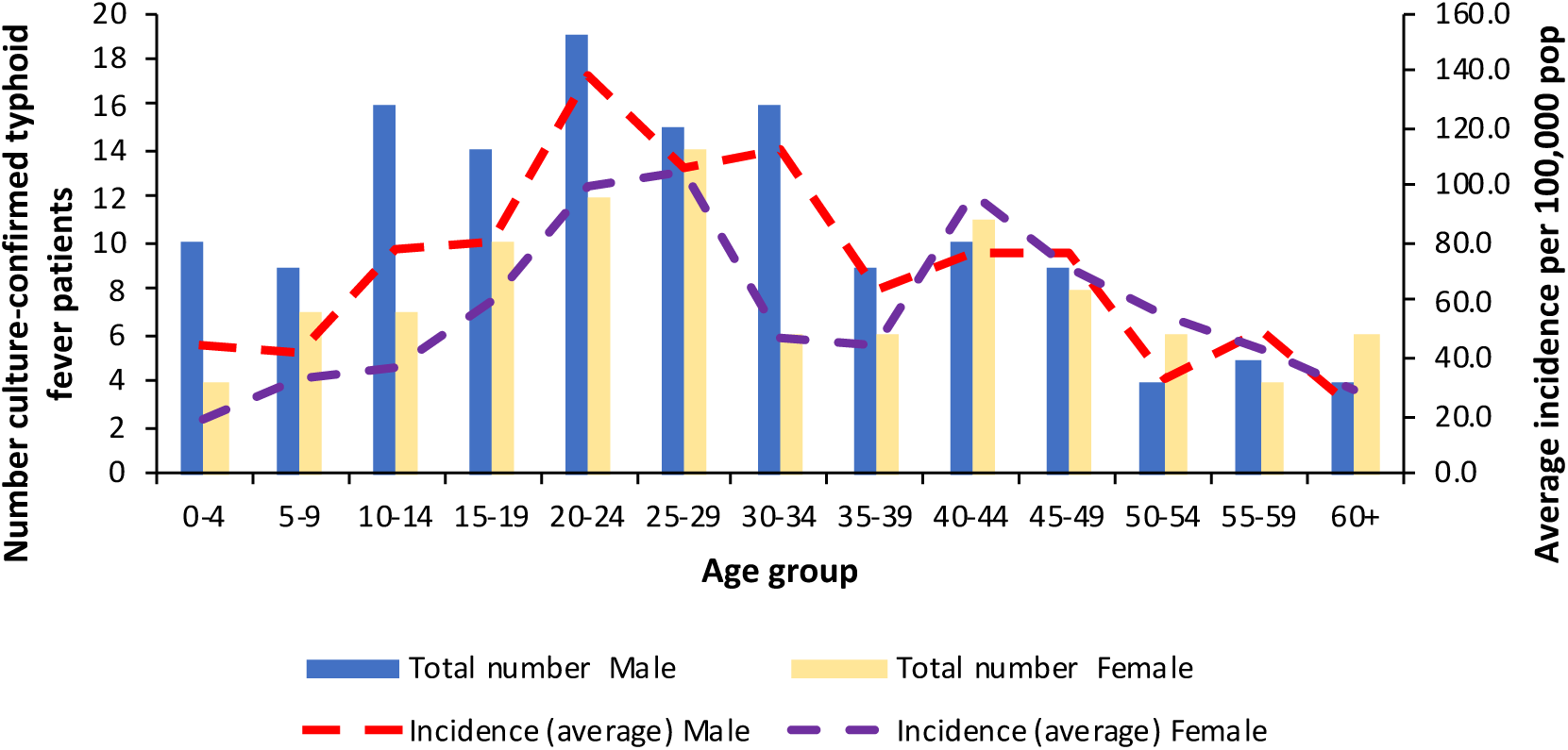
Number and incidence of culture-confirmed typhoid fever patients by age groups and gender in the Northern Division of Fiji between 2017-2019. Number of culture-confirmed cases are reflected by columns with average incidence per 100,000 population indicated by the dashed lines.

### Temporal characteristics of Northern Division typhoid patients

Typhoid fever patients were reported every month throughout the three-year study period, suggestive of endemic transmission. Fiji has two defined seasons: the wet season (associated with tropical cyclones) which runs from November to March, and the dry season (April to October). The distribution of typhoid fever patients did not show a clear seasonal trend (e.g., wet versus dry) during the 2017-2019 study period. The temporal distribution of typhoid fever patients was characterised by several peaks corresponding to outbreaks, as defined by the report of two or more patients with typhoid, in the same household or community, in a four-week period^21^. A total of 127 (51.6%) patients were epidemiologically linked to outbreak clusters in household or community settings. There were a total of 28 such clusters of which 13 were household clusters (n=30) and 15 community clusters (n=97).

### Genomic characteristics of Northern Division and Central Division *S*. Typhi isolates

A total of 240 *S*. Typhi isolates from 217 patients (86.1% of the total cases) from the Northern Division were collected and their genomes sequenced (Supplementary table 1). The remaining 13.9% of patients were excluded as stored samples could not be recovered. All strains in the Northern Division belonged to *S*. Typhi genotype 4.2, which fall into two lineages: 4.2.1 (30%, n= 73) and 4.2.2 (70%, n= 167) (Supplementary figure 1). There was some evidence of geographical restriction of lineages based on subdivision, where 4.2.1 was predominant throughout the three-year period in Bua subdivision, while in the remaining three subdivisions (Cakaudrove, Macuata and Taveuni) lineage 4.2.2 was prevalent in 2017 and 2018 before 4.2.1 became the more dominant by 2019 (Supplementary figure 1). This increase could be partly attributed to an unrecognised outbreak in Cakaudrove subdivision and subsequent transmissions to Macuata and Taveuni. There were no single or multi-drug resistant *S*. Typhi strains detected from the Northern Divisions by *in silico* analyses or phenotypic assay (Supplementary table 2).

To examine the relationship of typhoid strains within and between the two largely populated islands of Fiji (Vanua Levu island – Northern Division and Viti Levu island - Central Division), we sequenced the genomes of an additional 179 *S*. Typhi strains from the Central Division, collected contemporaneously i.e. between 2017-2019 (Supplementary table 1). Similar to the Northern Division, the majority (97.2%, 174/179) of the *S*. Typhi isolates were genotype 4.2, with lineages 4.2.1 and 4.2.2 accounting for 28.5% (n=51) and 68.7% (n=123), respectively. Two multi-drug resistant *S*. Typhi strains (genotype 4.3.1, H58) and three susceptible isolates (genotypes 3.5 and 3.5.4) were also identified in the Central Division in 2019 (Supplementary table 2).

Incorporating the 414 genotype 4.2 Central and Northern Divisions from 2017-2019 with our previous evolutionary framework of 367 Fijian *S*. Typhi genomes from (1981-2016)^12,22^ identified that the genotype 4.2 isolates rapidly expanded from 2008 onwards in Fiji despite the importation of regional genotypes (such as genotype 3.5 strains commonly associated with Samoa^11^) and the H58-like genotype 4.3.1 (Supplementary figure 2). These data indicate that sporadic importation of ‘global’ *S*. Typhi genotypes into Fiji occurs, yet such events have not resulted in displacement of the resident genotype 4.2 population over the last 15 years. This expanded temporal framework enabled refinement of the temporal dynamics of both genotype 4.2.1 and 4.2.2 in Fiji. For the 4.2.1 lineage, the mean evolutionary rate was estimated at 2.1 x 10^-7^ substitutions/site/year (95% highest posterior density (HPD) 1.4 x 10^-7^, 2.3 x 10^-7^) which is equivalent to 1 SNPs/genome/year (95% HPD 0.7, 1.4). This is slightly higher than estimates for 4.2.2 in Fiji (1.7 x 10^-7^ substitutions/site/year (95% HPD interval 1.5 x 10^-7^, 2.1 x 10^-7^). Our Maximum–clade credibility (MCC) tree shows the isolates in 4.2.1 lineage to be descendants of their most recent common ancestor (MRCA) in 1965 (95% HPD 1950.9 - 1976.2) (Supplementary figure 3). The isolates in the 4.2.2 lineage were most likely originated from their MRCA in 2005 (95% HPD 2002.7 - 2006.6) (Supplementary figure 3). The MCC tree also showed some evidence of a structured population with spatial clustering in 4.2.1 lineage. However, the 4.2.2 lineage did not exhibit the same population structure as 4.2.1. Similar to our previous findings^12^ the Skygrid analysis showed several contractions and overall reduction in effective population size of *S.* Typhi strains (Supplementary figure 4B/D).

### Phylogeographic and migration patterns of Fijian *S*. Typhi genotype 4.2

Phylogenetic analysis of the Fijian 4.2 lineage demonstrated substantial population structure in the tree indicating independent selection and spread of sub-lineages (Figure 2A). To further understand *S*. Typhi migration patterns within or between Central and Northern Divisions, we undertook a phylogeographical analysis of the 2017-2019 dataset based on the number of migration events (Markov jumps). The Northern Division is subdivided into 4 Sub-Divisions (Bua, Cakaudrove, Macauta and Taveuni). The predominant directionality of spread of both genotypes were largely between Sub-Divisions within the Northern Division, with Cakaudrove being a common migration source for genotype

**Figure 2.**
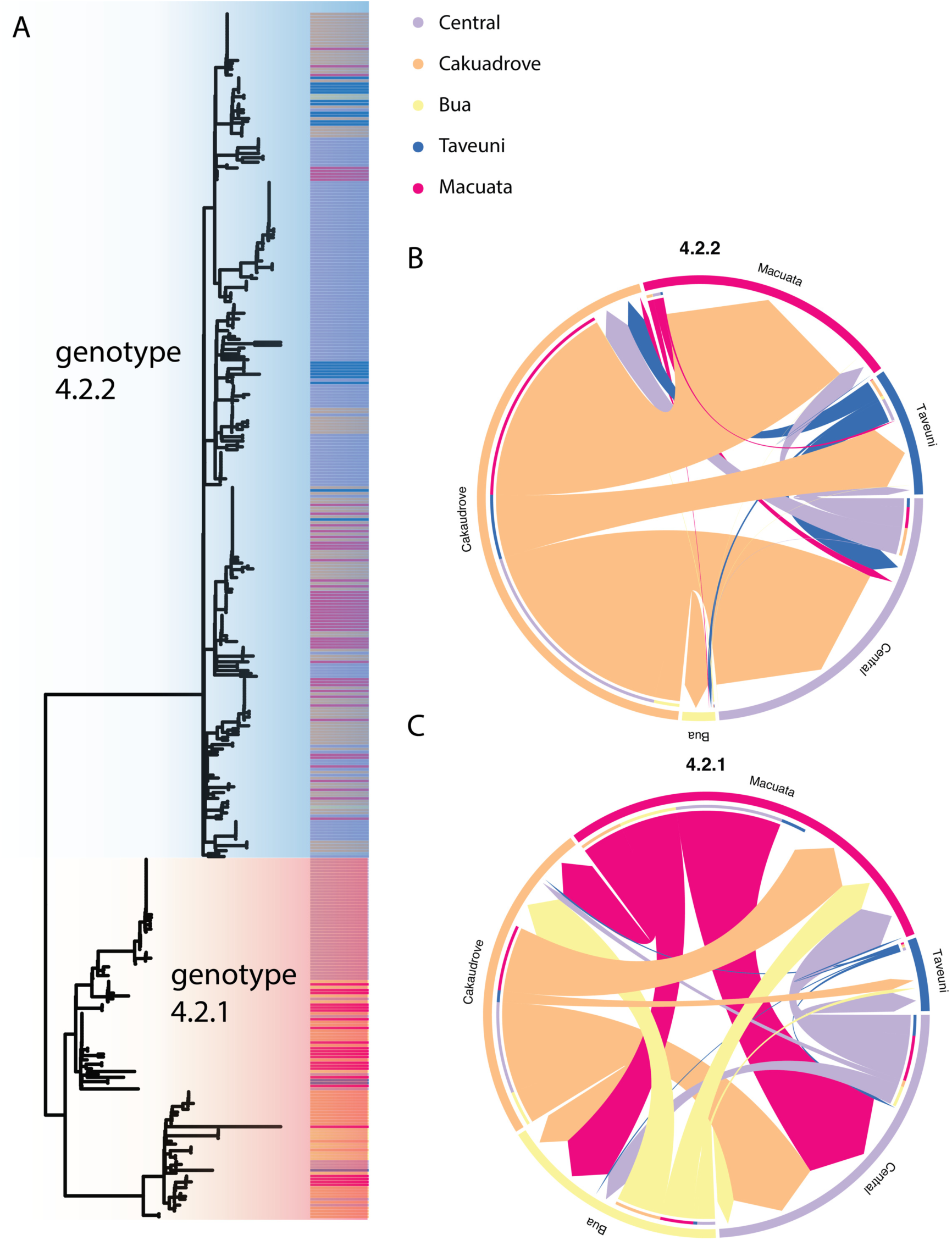
Phylogenetic structure of *S*. Typhi isolates in the Northern and Central Divisions, Fiji, between 2017-2019. **A)** Maximum likelihood tree of *S*. Typhi isolates from the Central Division (n=179) and the four subdivisions in the Northern Division (n=240) from 2017-2019 showing sub-division location (coloured by legend). **B)** and **C)** Chord diagram of migration patterns between sub-divisions inferred from mean counts of Markov jumps for genotype 4.2.2 (**B**) and 4.2.1 (**C**) populations (refer to Supplementary figures 5 and 6, Supplementary table 3).

4.2.2 and both Macuata/Cakaudrove for genotype 4.2.1 (Figure 2B and 2C respectively). This intra-divisional spread (within Central or Northern) was responsible for ∼95% of probable migration events in this dataset, with approximately 5% attributable to inter-division (between Central and Northern) migration. Of the inter-divisional migration events, on average, there were 2-fold and 4-fold more in the Northern-Central direction rather than Central-Northern, for genotype 4.2.1 and 4.2.2 respectively, although there were overlaps in 4.2.1 the migration counts (Supplementary table 3, Supplementary figure 6).

### Mutational profiling of the genotype 4.2 population

Despite the relatively low SNP mutation rate across the *S*. Typhi chromosome relative to other human pathogens^39^, recent studies propose that chronic carriage of *S.* Typhi can result in increased frequency of mutations in genes encoding membrane lipoproteins, transport/binding proteins, surface antigens and regulators ^18^. In particular, genes encoding for the Vi capsular polysaccharide have been shown to accumulate mutations in chronic carriage, and Vi genes have been shown to be missing in some isolate^18^. This finding prompted us to examine evidence of genetic selection across the Fijian genotype 4.2 dataset. A total of 1,265 mutational events (SNPs/small indels) relative to the Fiji genotype 4.2.2 reference strain ERL072973 (LT904777.2) were identified among the 782 genotype 4.2.1 and 4.2.2 *S*. Typhi isolates collected between 1981-2019 (Supplementary table 4). Two regions of the chromosome had elevated numbers of mutations: the *viaB* operon encoding for the Vi polysaccharide synthesis genes and the region containing an alternate sigma factor *rpoS*, a stress response regulator (Figure 3A). Overall, one third (34.1%, 257/782) of the *S*. Typhi strains had at least one *viaB* operon mutation (*tviA*-*tviE*) and 9.3% (73/782) had a *rpoS* mutation. *viaB* operon SNPs accounted for 2.8% of all mutational events (36/1,265) with 31 of these across two genes, *tviD* and *tviE* (Figure 3B & C). Of these mutations, 97% (30/31) encoded for protein coding changes (missense mutations), suggesting selective pressure may be acting to maintain amino acid altering mutations in these genes. *rpoS*, was also over-represented with 59 mutational events (4.7%), 31 encoding for missense mutations and 28 encoded for non-sense mutations (frameshifts/premature stop codons) (Figure 3B).

**Figure 3.**
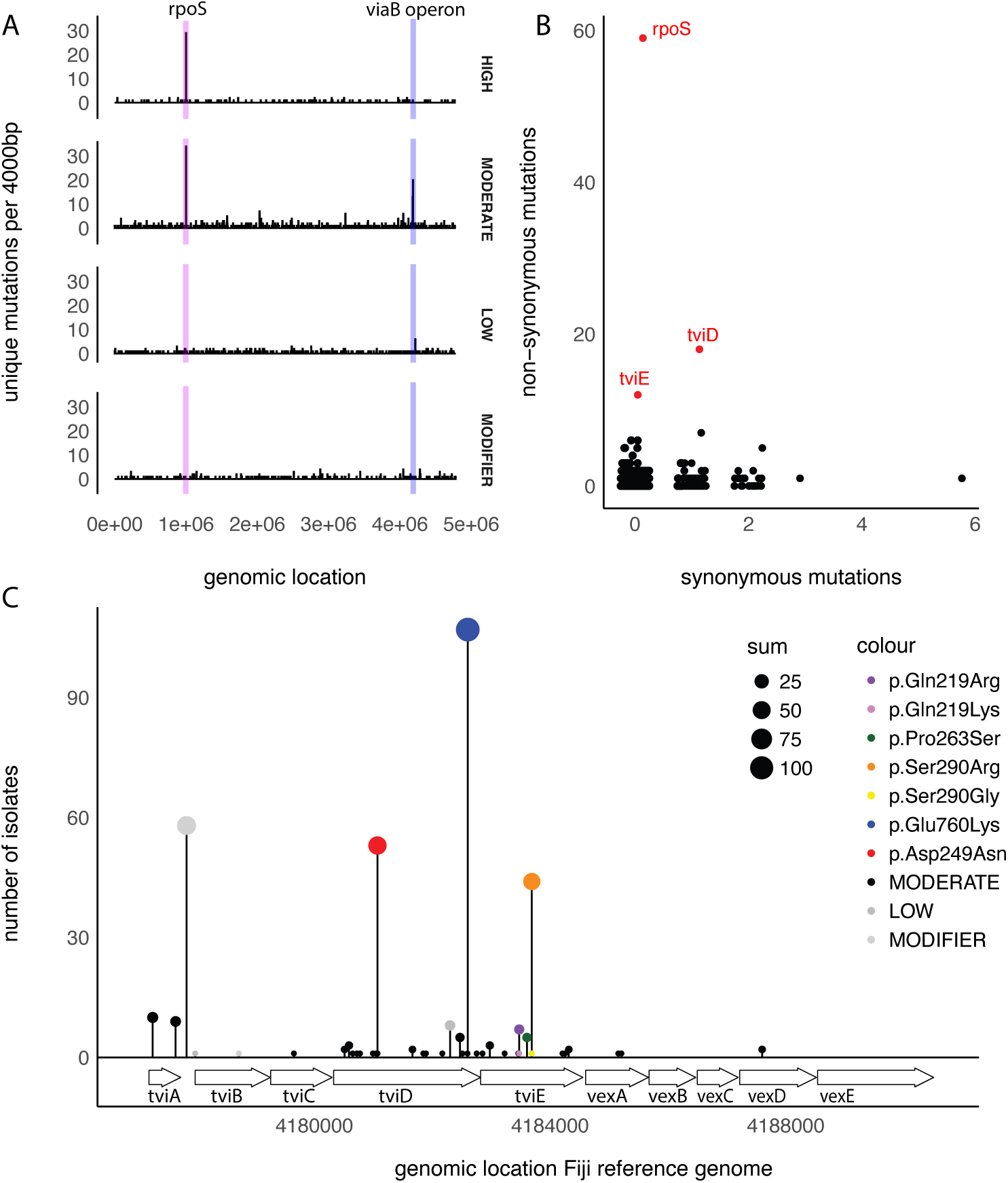
Preferential mutation of *viaB* operon and *rpoS* in the Fiji genotype 4.2 population. **A)** Histogram of 1,265 unique mutations (single nucleotide polymorphism and small indels) in the Fiji population with a 4,000bp binwidth. Two chromosomal regions encompassing the *rpoS* (purple) and *viaB* operon (blue) exhibit elevated mutation rates relative to the rest of the chromosome. Effect size of the mutation is faceted to show distinction between ‘high’ functional consequence (frameshift/truncation mutations), ‘moderate’ (non-synonymous, resulting in an amino acid change), ‘low’ (synonymous, no change in amino acid) and ‘modifier’ (intergenic region). **B)** Gene-based frequency of non-synonymous mutations (resulting in an amino acid change) versus synonymous mutations (non-protein changing). Genes containing over 10 independent mutations with ‘high’ or ‘moderate’ functional consequence are highlighted in red. *rpoS, tviD* and *tviE* genes have the highest number of unique non-synonymous mutations in the Fijian genotype 4.2 population. **C)** Lollipop plot of SNPs location within the *viaB* region. Size of dot and height of bar is correlated to the number of isolates. Non-synonymous SNPs that were deemed to persist in the population over an extended period or occurred multiple times independently are coloured as per legend. Both TviE Pro263Ser (green) and Gln219Arg* (purple) occur in both genotypes 4.2.1. and 4.2.2.

Despite this overrepresentation, the vast majority of mutations in the *viaB* operon and *rpoS* were in a limited number of isolates (3 or less). However, three mutations, Ser290Arg in TviE and Asp249Asn/Glu760Lys in TviD, that persisted throughout the study period (Supplementary figure 7). The isolates containing these mutations were geographically spread, suggesting that mutations in *tvi* genes can be maintained in the population through new infections (Supplementary figure 8). For example, the Ser290Arg TviE substitution was first evident in a 4.2.2 sub-lineage ∼2008, with cluster analyses revealing ongoing transmission of this *S*. Typhi clone mainly in the island of Taveuni from 2017 to 2019, without any documented community outbreak (Supplementary figure 7). In support of the hypothesis that selection is playing a role in the overrepresentation of the *tvi* mutations rather than evolutionary stochasticity, several mutations appeared in the *S*. Typhi population multiple times and independently. Alterations of Gln219Arg (n=7), Pro263Ser (n=7) in TviE were found to occur in both 4.2.1 and 4.2.2 lineages while Ser290Arg (n=45) occurred twice in sub-lineages of genotype 4.2.2 with a Ser290Gly (n=1) occurring in lineage 4.2.1 in 1984 (Supplementary figures 7 and 8).

### Global markers of *viaB* genetic adaptation

To determine if selection in the *viaB* operon and *rpoS* genes was a genotype 4.2-specific feature or if this mutational profile is represented across *S.* Typhi genotypes, a global dataset of 12,382 genomes, representing 86 genotypes (distinct genotyphi clusters) collected between 1916 to 2021^13^ was analysed. A total of 124,180 SNPs were observed across the 12,382 genomes relative to the archetypical *S*. Typhi reference genome CT18, with 36,432 SNPs (29%) present in more than one isolate. Of these SNPs that occurred at least twice, a similar pattern was observed as in the Fiji dataset where chromosomal regions encompassing the *rpoS* and the *viaB* operon contained the highest density of non-synonymous mutations (Supplementary figure 9). At a gene specific level, *rpoS, tviD* and *tviE* had amongst the largest ratio of non-synonymous to synonymous mutations and total count of non-synonymous mutations (Figure 4A). Each had many unique events within the population with a heavy bias towards non-synonymous mutations. *rpoS* contained 53 missense mutations, 23 nonsense mutations (a reduction relative to the Fiji dataset due to the exclusion of small indels) and 4 synonymous mutations. The genes *tviD* and *tviE* each contained several mutations with 1 nonsense, 128 missense and 9 synonymous mutations in *tviD* and 40 missense and 2 synonymous mutations in *tviE.* These were independent events of which some were found to occur multiple times in the population (Table 2 & Figure 4B, 4C).

**Figure 4.**
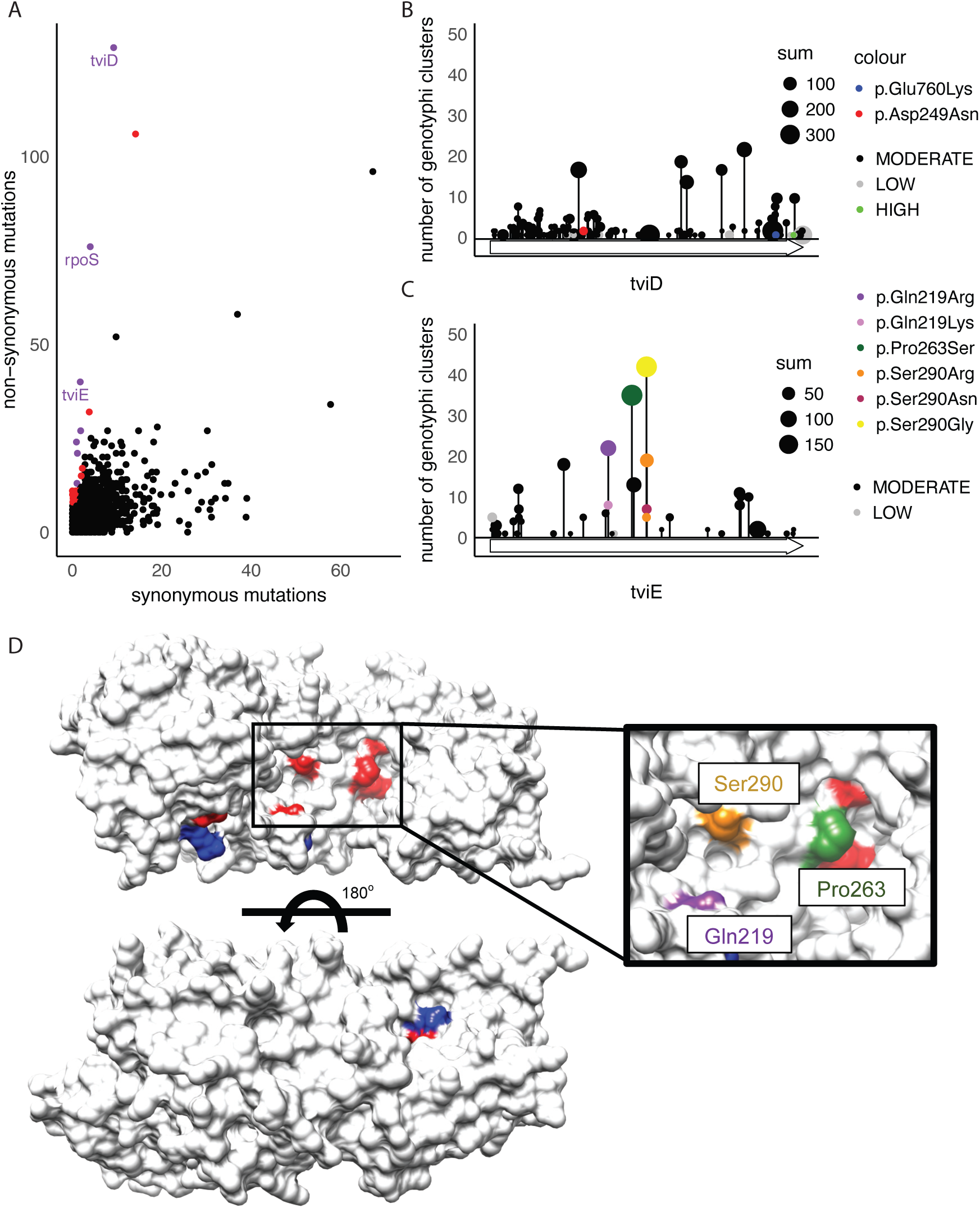
Independent occurrence of mutations in *tviD* and *tviE* genes in the global population of *S.* Typhi. Single nucleotide polymorphisms (SNPs) per gene were categorised into non-synonymous (predicted to have a predicted ‘high’ or ‘moderate’ consequence) and synonymous (defined as ‘low’ consequence). **A)** Gene-based frequency of non-synonymous mutations (resulting in an amino acid change) versus synonymous mutations (non-protein changing). Genes with a high ratio of non-synonymous mutations relative to synonymous mutations are highlighted in red (>7:1) and purple (>12:1). The *rpoS, tviD* and *tviE* genes have some of the highest number of unique non-synonymous SNPs in the global population. A single integrase gene (STY3193) with 65 non-synonymous and 207 synonymous mutations was excluded from this plot to enable better visualisation (refer to Supplementary figure 10 for complete plot). **B)** Lollipop plot of unique SNPs in the *tviD* and **C)** *tviE* genes. Size of dot is correlated to the number of isolates in the global population of 12,382 *S*. Typhi genomes containing the identical site-specific mutation and the height of the lollipop dots corresponds to the number of different genotyphi clusters where that mutation occurred. Specific amino acid mutations of interest from the Fiji population are coloured and highlighted (refer to legend) as per Figure 3C. The presence of two Ser290Arg dots is explained by two unique SNP events that both independently encode for a codon change required to switch to an Arginine. **D)** Mapping of non-synonymous mutations onto the predicted structure of TviE (uniprot Q04975) demonstrating that mutations are surface exposed and located on the same side of the protein predicted to contain an active site^17^ (Supplementary figure 11). Residues with missense mutations present in >10 genotyphi clusters highlighted in red, > 5 genotyphi clusters (95% quantile) in blue. Inset – the mutations highlighted in the Fiji dataset are coloured as per the lollipop plot (**C**) 90 degree rotations of Figure 4D are supplied as Supplementary figure 12.

**Table 2:** TviE mutations present in greater than 5 different genotyphi clusters (i.e. in the 95% quantile for number of unique clusters)

| Mutation | Isolates (%) | Clusters* (%) | Genotypi |
| --- | --- | --- | --- |
| p.His53Tyr | 23 (0.19) | 12 (14) | 2.0.2, 2.2, 2.3.2, 2.5, 3, 3.3.1, 3.3.2, 4.1.1, 4.2.3, 4.3.1.1, 4.3.1.2, 4.3.1.2.1 |
| p.Asp54Asn | 17 (0.14) | 7 (8) | 1.2.1, 2.5, 3, 3.3.2, 4.1.1, 4.3.1.1, 4.3.1.3.Bdq |
| p.Cys137Arg | 49 (0.39) | 18 (21) | 1.1, 2.1.7.1, 2.1.7.2, 2.2.2, 2.4, 2.5, 3.1.1, 3.2.2, 3.3.2, 4, 4.1, 4.2.2, 4.3.1, 4.3.1.1, 4.3.1.1.EA1, 4.3.1.2, 4.3.1.2.1, 4.3.1.3.Bdq |
| p.Thr214Lys | 8 (0.06) | 6 (7) | 1.2, 2, 3.1.1, 4.1, 4.3.1.1.EA1, 4.3.1.3.Bdq |
| p.Gln219Lys | 13 (0.1) | 8 (9) | 2.3.3, 4.1, 4.2.2, 4.3.1, 4.3.1.1, 4.3.1.1.EA1, 4.3.1.2, 4.3.1.2.1 |
| p.Gln219Arg | 90 (0.73) | 22 (25) | 2, 2.0.2, 2.2, 2.3.1, 2.3.2, 2.3.4, 2.5, 3.1, 3.1.1, 3.2.2, 3.3.2.Bd2, 3.4, 3.5, 4.1, 4.1.1, 4.2.2, 4.3.1, 4.3.1.1, 4.3.1.1.EA1, 4.3.1.1.P1, 4.3.1.2, 4.3.1.2.1 |
| p.Pro263Ser | 194 (1.56) | 35 (41) | 2, 2.1.4, 2.2, 2.3.2, 2.3.3, 2.3.5, 2.4, 2.4.1, 2.5, 2.5.2, 3, 3.0.1, 3.1, 3.1.1, 3.2.1, 3.2.2, 3.3, 3.3.1, 3.3.2, 3.3.2.Bd2, 3.4, 3.5, 3.5.4.1, 3.5.4.3, 4, 4.1, 4.1.1, 4.3.1, 4.3.1.1, 4.3.1.1.EA1, 4.3.1.1.P1, 4.3.1.2, 4.3.1.2.1, 4.3.1.2.EA2, 4.3.1.3.Bdq |
| p.Lys266Asn | 79 (0.64) | 13 (15) | 0, 0.0.1, 0.0.2, 0.0.3, 0.1, 0.1.1, 0.1.2, 0.1.3, 1.1, 1.1.1, 1.1.2, 1.1.3, 1.1.4 |
| p.Ser290Gly | 188 (1.52) | 42 (48) | 0, 0.0.1, 0.0.2, 0.0.3, 0.1, 0.1.1, 0.1.2, 0.1.3, 1.1, 1.1.1, 1.1.2, 1.1.3, 1.1.4, 2, 2.1.7.1, 2.1.7.2, 2.2, 2.2.2, 2.3.2, 2.3.3, 2.3.4, 2.4, 2.4.1, 2.5, 3, 3.1.1, 3.2.1, 3.2.2, 3.3, 3.3.1, 3.3.2, 3.5, 3.5.4.3, 4.1, 4.1.1, 4.2, 4.2.1, 4.3.1, 4.3.1.1, 4.3.1.1.EA1, 4.3.1.2, 4.3.1.2.1 |
| p.Ser290Asn | 19 (0.15) | 7 (8) | 2.1.7.1, 3.2.1, 3.2.2, 3.3.1, 3.4, 4.3.1.1.EA1, 4.3.1.2 |
| p.Ser290Arg | 70 (0.56) | 19 (22) | 2, 2.1.7.1, 2.2.4, 2.3.3, 2.5, 3, 3.2.2, 3.3, 3.3.2, 3.3.2.Bd2, 3.5, 3.5.4.3, 4.1, 4.2.2, 4.3.1, 4.3.1.1, 4.3.1.1.EA1, 4.3.1.2, 4.3.1.2.1 |
| p.Ala462Thr | 34 (0.27) | 11 (13) | 2, 2.2.2, 2.3.5, 3, 3.1.1, 3.2.2, 3.3.2, 3.5, 4.1, 4.3.1.1, 4.3.1.2 |
| p.Ala462Val | 23 (0.19) | 8 (9) | 2, 2.0.2, 2.1, 2.2.1, 3, 3.2.2, 4.1.1, 4.3.1.1 |
| p.Arg464His | 11 (0.09) | 10 (12) | 2.1.7.1, 2.2, 3.1.1, 3.2.2, 3.3.1, 3.3.2, 3.5, 4.3.1, 4.3.1.2, 4.3.1.2.1 |
| p.Phe479Leu | 16 (0.13) | 10 (12) | 2, 2.2, 2.3.2, 3, 3.1, 3.2.2, 3.3.1, 4.3.1.1.EA1, 4.3.1.2, 4.3.1.2.1 |

\*Maximum number of genotypi clusters is 86

While there were over 100 independent non-synonymous SNPs in *tviD* in the 12,382 isolates, few (15/138, ∼10%) were found in greater than 5 genotyphi clusters. Furthermore, the specific *tviD* non-synonymous mutations that were observed in large numbers of isolates in the Fiji genotype 4.2 population did not appear to be highly prevalent in the global population. For example, the Asp249Asn and Glu760Lys mutations which appear to have been maintained in the Fiji genotype 4.2 dataset (Figure 3B) were exclusive to this genotype. In contrast, 15 of 42 *tviE* mutations (35%) were in greater than 5 genotyphi clusters. This included all 3 *tviE* mutations that were found to occur independently in genotype 4.2.1 and 4.2.2. Gln219Arg occurred in 22 clusters (90 isolates including 29 isolates in genotyphi 2.3.1), while a related mutation Gln219Lys occurred in 8 genotyphi clusters (n=13). Pro263Ser occurred in 35 unique genotyphi clusters (194 isolates including a cluster of 10 genotyphi 2.3.5). Ser290Arg occurred in 19 unique genotyphi clusters (70 isolates) with 2 independent mutations causing this change in 5 genotyphi clusters. The same serine 290 residue was also mutated to Glycine and Asparagine in the global population. Ser290Gly was present in 42 genotyphi clusters (188 isolates and is omnipresent in serotype 0.X and serotype 1.1.X) and Ser290Asn present in 7 clusters (19 isolates) (Figure 4C, Table 2; Supplementary table 5). This observation suggests that there have been multiple independent selections of these mutations occurring across different genetic backgrounds and in geographic locations. In addition to these mutations, an additional 4 mutations were observed in greater than 10 genotyphi clusters (Cys137Arg, Lys266Asn, His53Tyr, Ala462His) and 4 sites in greater than 5 genotyphi clusters (Arg464His, Phe479Leu, Asp54Asn/Tyr, Thr214Lys). Despite this increased population-level of SNP frequency in the *tvi* locus, individual isolates tend to only have a small number of missense mutations in *tviD* and *tviE* with no isolate containing more than 4 mutations relative to the most common sequence and only 2 isolates containing more than 4 mutations across both genes. This may indicate a possible fitness cost associated with accumulation of *tvi* mutations.

To better understand the likely effect of these mutations, we mapped the mutated residues that were present in greater than 5 genotyphi clusters (Table 2) to a putative Alphafold structure of TviE. The location of the evolutionary independent mutations localised to the same surface region of the TviE protein (Figure 4D, Supplementary figure 12), which are adjacent to predicted active site residues defined in Wear et al.^17^.

## Discussion

Typhoid fever is a health concern in many under-resourced settings around the globe, such as the South Pacific Island nation of Fiji, where elevated rates of infection constitute a substantial public health burden^3–4^. To better define the evolutionary dynamics of *S*. Typhi across the two main populated isolates of Fiji (Viti Levu and Vanua Levu), we carried out a genomic epidemiology and phylogeographic study of the *S*. Typhi isolates using representative samples (85% of the total reported cases) from the Central and Northern Divisions between 2017-2019.

Analysis of these genomes in the context of previous sequences from Fiji (1980-2016)^12,22^ revealed a shift in *S.* Typhi population structure over the past 40 years. Between 1980 and mid 2000s, a wide range of genotypes were sequenced. However, from 2008 onwards typhoid fever in Fiji has primarily been caused by a single genotype (4.2) with two lineages, 4.2.1 and 4.2.2, circulating concurrently in the country^12,22^. A review of publicly available genomic data (>12,000 isolates) showed the 4.2.1 and 4.2.2 lineages largely have only been detected outside of Fiji in travel-associated cases with only a single isolate being identified in New Zealand in 2019 with no travel history^41^. These findings support the theory that the two lineages are endemic in Fiji^12,22^. Our analysis showed sporadic importation of 4.3.1 (MDR genotype) and 3.5 genotypes with no evidence of circulation of these strains in the community, or ongoing in-country transmission. We can conclude that typhoid fever in Fiji most likely occurs through expansion, selection and regional transmission of ‘local’ *S*. Typhi strains (genotype 4.2) over time, rather than through importation and replacement with ‘new’ extant strains. These findings are commensurate with recent studies of other Pacific Island countries that *S*. Typhi genotypes appear endemic in nature^10,11^. These data demonstrate that while foreign importations of multi-drug resistant clones (such as genotype 4.3.1) are evident, they have not displaced the endemic genotypes in these regions to date. This observation is in contrast to the rapid spread of MDR genotype 4.3.1 and associated sub-lineages which replaced the local *S*. Typhi population in Africa and throughout Asia^22,41–44^. The *S*. Typhi population in Fiji is recognised for low levels of antimicrobial resistance^5–7^, which is also the case in this study. This may in part be due to the restricted antimicrobial prescribing practices in Fiji, which are accessed by prescription only, and the supply of key drugs like ciprofloxacin is strictly regulated. Thus, ongoing antimicrobial stewardship practices may be a contributing factor for the absence of substantial antimicrobial resistance among the endemic *S*. Typhi population.

Our data revealed a dynamic an ongoing interplay of two endemic *S.* Typhi sub-lineages. The 4.2.1 lineage originated from its MRCA in 1965 and has persisted in Fiji for over 50 years. This lineage is evolving at slightly faster rate and exhibits substantial genetic diversity compared to the 4.2.2 lineage and other global genotypes^10,39,44,45^. Clonal expansion of 4.2.1 lineage attributed to large scale community outbreaks or ongoing transmission was also more pronounced from 2015 onwards. Ongoing genomic surveillance is needed to monitor the evolution of this lineage as it can result in the creation of separate sub-lineages.

Our genomic analysis of *S*. Typhi genotype 4.2 revealed a disproportionately higher density of non-synonymous SNPs in *viaB* operon. The *viaB* operon, located in SPI-7, is a genomic island approximately 134 kb in size^46^ that has 10 genes (*tviA* to *E* and *vexA* to *E*) which are involved in the synthesis, export and anchoring of the Vi polysaccharide^15,16,47^. The surface exposed Vi capsular polysaccharide is an important virulence determinant^48^. We found 30 distinct non-synonymous point mutations among the Fijian *S*. Typhi isolates with almost one third of isolates have at least one *viaB* SNP. Our findings also confirmed the persistence and clonal expansion of *S*. Typhi strains with *viaB* point mutations (notably in *tviD and tviE* genes) in both endemic lineages and across the two main islands. We also demonstrated independent acquisition of non-synonymous point mutations in TviE (at codon Gln219Arg and Pro263Ser) across *S*. Typhi genotype 4.2 lineages in different temporal and geographic locations. *S*. Typhi strains with multiple *viaB* mutations were also found in both acute and carrier state isolates. The above findings indicate selective pressure and potential adaptive selection among endemic *S*. Typhi lineages in Fiji.

With this in mind, an analysis of the prevalence of Vi mutations in a globally diverse set of sequences was also undertaken. Globally, mutations in *viaB* operon are considered uncommon^18,19,45^. Recent studies reported non synonymous SNPs in *tviB, tviD* and *tviE* genes in *S.* Typhi strain from Nepal^18^ and Kenya^45^ yet these studies detected small numbers of mutations (fewer than seven isolates) mainly among isolates from carriers. To our knowledge, the occurrence of multiple *viaB* mutations occurring between *S.* Typhi strains cultured from acutely sick patients, as seen in our surveillance study, have not be reported in the literature to date^19^. We observed common *tviD* and *tviE* mutations arising independently in multiple evolutionary distinct lineages globally and, in a few cases, located in transmitting populations of acute patients rather than in chronic carriers.

Several other genes showed evidence of increased non-synonymous mutation rate in this study. The *rpoS* gene was shown to have many independent mutations leading to premature truncation and loss of function. This phenomenon has previously been reported with 15 of 41 *S.* Typhi isolates containing a defective *rpoS* in one study^49^ and *rpoS* mutations over-represented in cases of gall bladder chronic carriage^18^. In contrast to the *viaB* mutations reported here, there was limited evidence of clonal expansion for *rpoS* mutants in the Fiji dataset with a maximum of two related isolates containing the same mutation. This trend was also observed in the global analyses where of all independent *rpoS* mutations, only 5 occurred in more than 10 isolates, many of which were stochastic in nature. One exception was a stop codon (nucleotide 2,915,627 in CT18, p.Trp148*) being present in multiple related isolates (n=5, genotyphi 4.1) from 2007-2010 in Indonesia, suggesting transmission of *rpoS* mutants is possible over short-time scales yet are unlikely to be maintained in the population. The total number of rpoS frameshifts reported in our global analyses is likely an under-representation given that small indels resulting in *rpoS* frameshifts mutations were excluded from the analysis. Other genes showed an elevated rate of non-synonymous mutations in this study, including the genes encoding the two-component regulatory system YehUT (STY2388/2389), of which STY2389 was previously identified in the study of Gall bladder chronic carriage^18^ This two-component system has been shown to regulate the carbon starvation protein CtsA (44). Further investigation of the impact of these mutations was outside of the scope of this study yet a full list of mutations and their associated genotyphi association is provided (Supplementary table 5).

The overall impact of the *viaB* mutations on the Vi-capsular structure and other phenotypic characteristics are presently unknown. All *S.* Typhi isolates in this study, including those with *viaB* SNPs were found to be Vi-antigen positive as determined by slide agglutination with Vi typing antisera. The altered residues therefore are not likely to convey a loss of function. While the binding site of the sugar-nucleoside ligand (UDP-GalNAcA) in TviE is unknown, recent studies identified an EX_7_E motif in TviE (13), predicted to be involved in sugar nucleoside binding. Mutations of either residues E483 or E491 in this motif within a TviE expression construct was unable to produce Vi antigen, suggesting that these residues are important for the polymerisation of the nucleoside sugar. While the mutations we observed in this study were adjacent to this motif, these two residues are not mutated in any of the isolates in the global dataset. While single point mutations of catalytically important residues are capable of greatly altering the enzymatic function of TviE^17^, there are other potential roles that mutations could play. Wear *et. al*^17^ predicted a protein-protein interface between TviD and the non-catalytic elements of TviE which may enable the enzymes to act in concert. If the mutations we observed were involved in this interface, the efficiency of the O-acetylation step rather than the polymerisation may be affected. Without a definitive ligand-bound structure, the effect of the mutations we identified in the global population of *S.* Typhi remains to be elucidated. Further research is necessary to investigate the risk factors associated with high rate of mutations in genes clusters involved in surface polysaccharide synthesis and transport among the endemic *S*. Typhi lineages in Fiji and determine phenotypic impacts on the Vi-capsule structure and function.

Vi capsular polysaccharide is an immunogenic antigen and is the target for purified subunit as well as the recently WHO prequalified conjugate vaccine that are being used in many endemic countries for typhoid fever control^50,51,52,53^. Between 2010 and 2017, several targeted mass vaccination campaigns using Vi-polysaccharide vaccine have been used widely in Fiji (including the study sites) to control community-based outbreaks^6,54^. It has been postulated that mass vaccination with Vi-containing vaccine could increase selection pressure on *viaB* operon which could lead to loss of Vi capsule^55,56^. However, this was not the case in this study. Though SNPs in *viaB* operon may be indicators of this potential threat; all the isolates we examined were still Vi positive. As part of the medium-term typhoid control strategy, mass vaccination with Typbar TCV begun in the Northern Division in 2023. Ongoing phenotypic and genomic characterisation of Vi capsular polysaccharides in *S*. Typhi isolates before and after mass vaccination is warranted to assess population impacts.

There are other potential driving forces for selection of mutations in the *viaB* operon other than host immunity or vaccination. Previous studies have shown that deletion of *tviD* results in approximately 10-fold increase in survival in water^57^, though the mechanism is unknown. Further, the existence of strain specific Vi typing phage suggests that modification of this surface marker may enable escape from predation of some naturally occurring lytic phage, producing a selective pressure for variation in the population^58^. Understanding the functional effects and immunological consequence of *tvi* operon mutations on Vi phenotypes would aid in assessing the biological and therapeutic impact of recurring *tvi* mutations globally.

## Acknowledgments

The authors would like to thank the staff of Labasa hospital, Northern Health, Colonial War Memorial Hospital staff and Fiji Centre for Communicable Diseases Control who supported in data collection, facilitated sample collection and laboratory testing. We would also like to thank Ms Helen Thomson (Murdoch Children’s Research Institute, Melbourne, Australia) for providing administrative support and Ms Silivia Matanitobua, (Fiji Centre for Communicable Diseases Control) for facilitating isolates shipment. We acknowledge the support of staff at the Microbiological Diagnostic Unit Public Health Laboratory.

## Authors Contributions

Authors contribution: AGS, RAS, KM, and MRD conceived the study and designed the study protocol. AGS and VR conducted epidemiological data collection. MM, TS and OC facilitated local approvals and assisted in data collection OC, and VS performed laboratory procedures. MV, and BP facilitated genome sequencing of the isolates. AGS, AH, WW, JAL, RAS, SD and MRD performed data analysis. AGS, AH, WW, RAS and MRD wrote the manuscript. AGS, AH, WW, MM, TS, OC, VS, VR, NW, JAL, DH, MV, AJ, BPH, SD, KM, RAS and MRD read and approved the final manuscript.

